# High-Throughput Screening for Myelination Promoting Compounds Using Human Stem Cell-derived Oligodendrocyte Progenitor Cells Identifies Novel Targets

**DOI:** 10.1101/2022.01.18.476755

**Authors:** Weifeng Li, Cynthia Berlinicke, Yinyin Huang, Weixiang Fang, Celeste Chen, Felipe Takaesu, Xiaoli Chang, Yukan Duan, Calvin Chang, Hai-Quan Mao, Guoqing Sheng, Stefanie Giera, James C. Dodge, Hongkai Ji, Stephen Madden, Donald J. Zack, Xitiz Chamling

## Abstract

Promoting myelination capacity of endogenous oligodendrocyte precursor cells (OPCs) is a promising therapeutic approach for central nervous system demyelinating disorders such as Multiple Sclerosis (MS). To aid in the discovery of myelination promoting compounds, we generated an advanced, genome engineered, human pluripotent stem cell (hPSC) line that consist of three reporters (identification-and-purification tag, GFP, and secreted NanoLuc) driven by the endogenous *PDGFRα*, *PLP1* and *MBP* genes, respectively. Based upon this line, we established a high-throughput drug screening platform and performed a small molecule screen with 2500 bioactive molecules. In addition to a number of previously known pathways, our screening effort identified new pathways whose inhibition enhance oligodendrocyte maturation and myelination. Although further genetic and molecular validation is required, the identified inhibitors could potentially be repurposed to develop remyelination therapy for MS and other demyelinating disorders.

## INTRODUCTION

Multiple Sclerosis (MS) is an immune-mediated demyelinating disease in which the immune system attacks and degrades myelin sheaths of neuronal axons in the central nervous system (CNS) (*1*). Damage and loss of a neuron’s myelin sheath can cause apoptosis of the myelin producing oligodendrocyte cells, axonal damage and neuronal cell death, which ultimately can lead to neurological disabilities (*2*). All currently approved therapies for MS act by modulating the patient’s immune response. Such therapies can help to slow down the progression of the disease, but do not directly promote the remyelination that is needed to improve the long-term function and survival of damaged neurons. A number of studies have shown that following myelin damage, oligodendrocytes progenitor cells (OPCs) and neural precursor cells (NPCs) migrate towards the site of injury, where they could differentiate into mature oligodendrocytes that are capable of remyelinating the demyelinated axons (*3*, *4*). However, endogenous progenitors around a CNS lesion appear to be limited in both their mitotic competence and differentiation potential, and they progressively lose their remyelination capacity with aging (*5*, *6*). If we could use biologically active molecules to increase the ability of OPCs to differentiate and become “myelinogenic”, this could have great therapeutic value for MS and other demyelinating diseases.

Drug screening to identify molecules that promote OL maturation and myelination has been initiated using primary rodent OPCs (*7*, *8*) and mouse ES-derived OPCs (*9*). The advent of human pluripotent stem cells (hPSC)-derived OPCs and OLs now make it possible to perform such drug screening using human cells. However, to our knowledge, no platforms that utilize human OPCs to screen for myelin promoting compounds are available yet. Use of the hPSC-derived, physiologically relevant cells for drug screening to identify molecules that promote remyelination is limited by the difficulty in obtaining large quantities of pure, disease relevant cell-types such as the OPCs (*10*). In addition, the majority of the OL maturation assays either use immunofluorescence analysis to measure the amount of MBP, or use image-based morphological analysis to detect cells that resemble oligodendrocytes (*7*–*9*, *11*, *12*). Although such image-based, high-content screening (HCS) assays have advantages, unlike in high-throughput screening (HTS), smaller differences are easily missed in the image-based analyses and they are also less scalable. Here, we used CRISPR/Cas9-based genome editing to generate a triple reporter hESC line where an identification-and-purification (IAP) tag (*13*), GFP, and secreted Nanoluciferase (secNluc) reporters were engineered to be driven by the endogenous *PDGFRα*, *PLP1* and *MBP* genes, respectively. Once the reporter hESC is differentiated, the PDGFRα expressing OPCs can be purified using the IAP tag (*13*, *14*), and the time and efficiency of oligodendrocyte maturation can be quantified via the expression of GFP and secreted Nluc. These qualities enabled us to use the reporter cell line to develop a highly sensitive and scalable screening platform for discovery of myelination promoting compounds that: 1) uses purified human OPCs, 2) allows time-course assays to follow real-time expression of oligodendrocyte/ myelination markers, and 3) makes it possible to perform Nluc-based HTS and image-based HCS using the same cell cultures.

Using the hOPCs derived from this reporter system, we screened 2500 bioactive molecules and identified several molecules that function at nanomolar doses to enhance maturation of oligodendrocytes from OPCs. At least two compounds that have not been previously reported to improve OL differentiation and enhance myelination were identified by this screen. Image-based analysis and electrospun nanofiber-based *in vitro* myelination assay further validated the effect of these compounds in hOPCs-hOL maturation. We also validated the efficacy of these compounds in a different hPSC-OPC reporter cell line. The identified pathways provide promising new potential targets for the development of remyelination-based therapies for the demyelinating diseases.

## RESULTS

### Generation of an advanced hESC reporter system

We previously generated a hESC reporter cell line (PDTT) that allows identification and purification of hESC-derived, PDGFRa expressing oligodendrocyte lineage cells (OLLCs)(*13*). Here, we added two more reporter sequences to the PDTT line and engineered an advanced reporter system that is suitable for HTS as well as HCS. CRISPR-Cas9-based genome editing was used to knock-in super-fold GFP (sfGFP) and secreted Nluc reporter sequences before the stop codons of the *PLP1* and *MBP* genes, respectively (Figure 1a). The PLP1-sfGFP construct generates a fusion reporter while the MBP-P2A-secNluc construct leads to two separate proteins (MBP and secNLuc) due to the presence of a self-cleaving P2A peptide.

**Figure 1.**
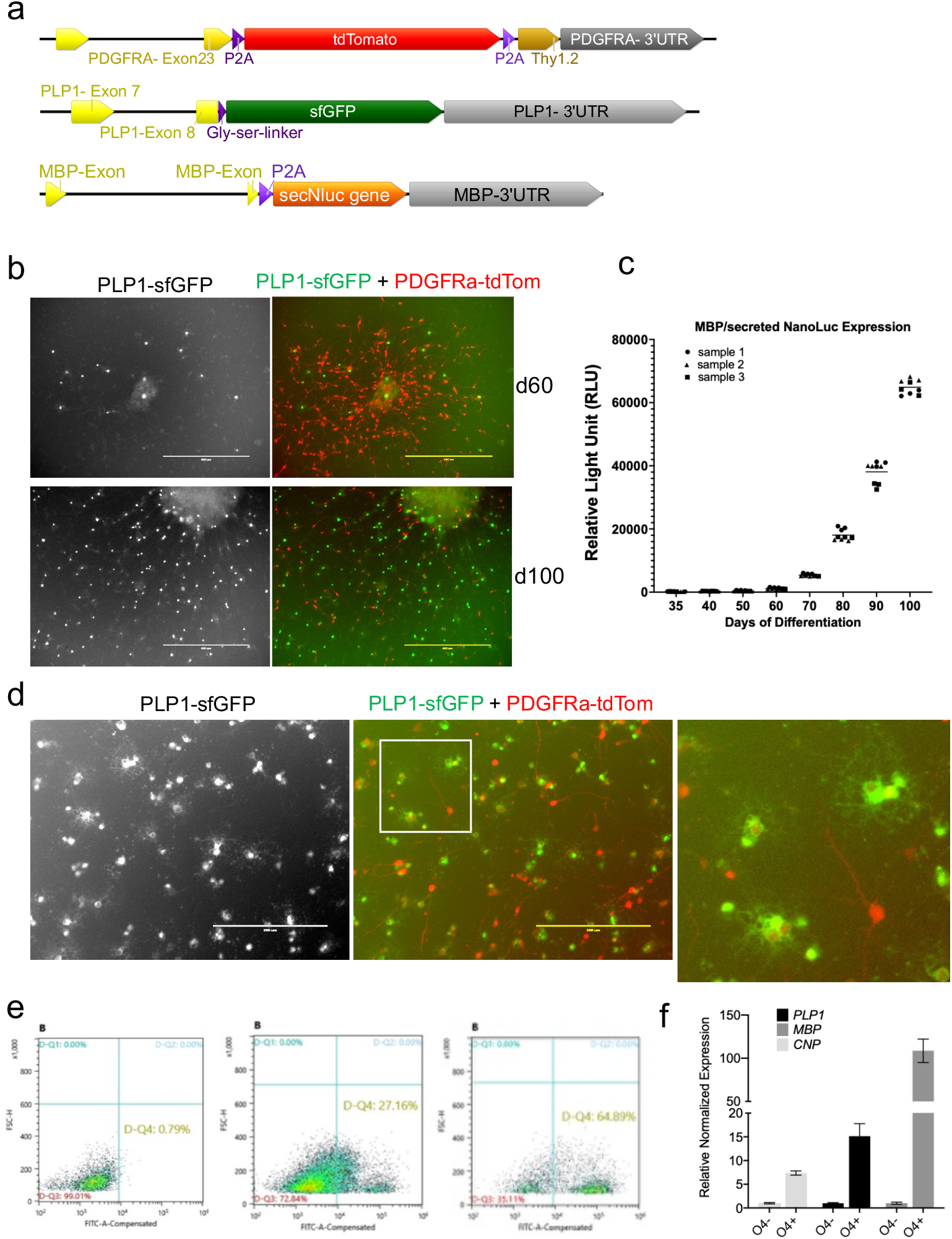
Generation of and validation of an advanced hES reporter system. **a)** Schematics of the endogenous *PDGFRA*, *PLP1* and *MBP* locus of the Ptt-P1-MsnL reporter line after CRISPR-Cas9-based genetic knock-in of reporter sequences. **b)** Expression of the tdTomato and sfGFP fluorescent reporters in the differentiating oligodendrocyte cultures. **c)** secNluc activity in the cell culture medium of differentiating PtT-P1-MsNL reporter line measured with Nano-Glo assay. Culture media from different days of differentiation (x-axis) were removed for the assay. Increase in Nano-Luc activity(y-axis), which corresponds to MBP expression increases as cells mature. **d)** Higher magnification of the reporter expression on d100 culture shown in b). GFP+ processes and branches resembling oligodendrocytes morphology are magnified in the inset. **e)** MACS-based enrichment of the GFP+ cells using O4 antigen microbeads. 65% PLP1-GFP+ cells were achieved from d105 cells. **f)** qPCR shows enrichment of OL markers in the O4+ cells compared to the O4-cells.

Prior to knocking-in the P2A-secNluc reporter into the *MBP* locus, we wanted to confirm that the secNLuc protein product, following its P2A-mediated separation, is successfully secreted into the extracellular culture media and retains enzymatic activity. Therefore, we transfected a CMV-tdTomato-P2A-secNluc plasmid into HEK293 cells and confirmed that the culture media of the transfected cells has Nluc activity (Figure S1a-c).

The final reporter system that we generated (PTt-P1-MsNL) thus incorporates three reporters: 1) PDGFRa-P2A-tdTomato-P2A-Thy1.2, in which, upon *PDGFRα* expression, tdTomato as well as Thy1.2 are produced. tdTomato localizes to the cytoplasm whereas Thy1.2 migrates to the cell surface, allowing the PDGFRa expressing cells to be immunopurified via Thy1.2 antibody conjugated magnetic microbeads (*13*); 2) PLP1-sfGFP, where expression of sfGFP is driven by *PLP1*; and 3) MBP-P2A-secNLuc, where NLuc protein product, which represents *MBP* expression, is secreted into the cell culture media (Figures 1a and S1d-f).

### Validation of the reporter cell line

In the PTt-P1-MsNL reporter line, similar to the PDTT cell line, PDGFRα-tdTom+ cells appear around day 45 of differentiation in our culture system(*13*). PLP1-GFP+ cells are visible as early as day 60. As the culture matures, less tdTomato+ cells and more GFP expressing cells are detected, and more MBP/sNLuc is also produced (Figures 1c and S2). The GFP+ cells have processes and morphology that resembles oligodendrocyte cells (Figure 1d). Moreover, the majority of GFP+ cells also express low levels of tdTomato, which indicates that these cells originated from PDGFRa expressing progenitor cells (Figures 1d and S2). We also show that an antibody against O4, an antigen specific to pre-oligodendrocyte cells, can be used to enrich PLP1-GFP+ cells from the differentiating OLLC culture (Figures 1e-f and S3c). Essentially, in this advanced reporter cell-line, PDGFRa+ OLLCs can be temporally purified (*13*), mature OLs can be identified and enriched for using the expression of PLP1-sfGFP, and the NLuc activity, which represents MBP expression, can be quantitated over time using a small aliquot of cell culture media.

### Optimization of the high throughput screening platform

An advantage of our reporter system is that it does not require fixation or lysis for assessment, which allows us to perform time-course assays to follow real-time expression of *PLP1* and *MBP*, oligodendrocyte /myelination marker genes. Since our main goal in generating the reporter system was to perform HTS, we went on to optimize our assay to fit a 384-well plate format. The seeding density, NanoGlo reagent volume, culture media volumes, and the optimal length of the assay were optimized in preliminary studies (Figures S4a-b) (see the methods section). Tasin-1, an inhibitor of Emopamil binding protein (EBP) and a known inducer of OL differentiation and myelination(*11*), was used as a positive control and DMSO was used as negative control for optimization of our assay (Figures 2a, S4c-e).

**Figure 2.**
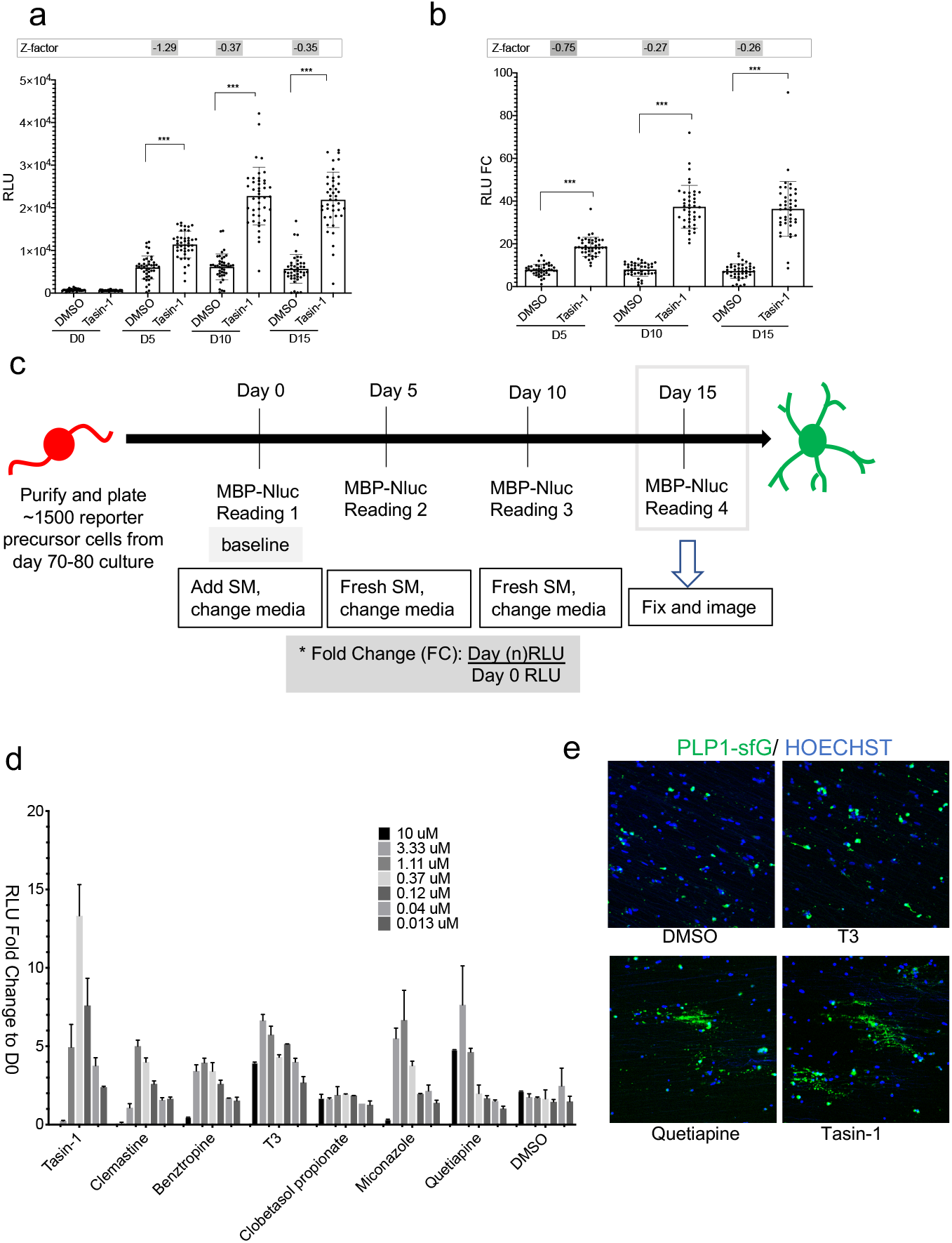
Drug screening assay optimization and validation. **a-b)** Effect of 250nM Tasin-1 in MBP-NanoLuc expression at days 5, 10, and 15 presented as **a)** raw RLU value and **b)** RLU fold change (FC). The Z-factor score for FC is better than the raw RLU. **c)** Timeline of the drug screening assay used for our reporter cell system. **d)** Validation of our drug screening assay using the compounds known to enhance oligodendrocyte differentiation. **e)** purified OPCs were plated on a plate containing electrospun nanofibler in the presence of DMSO or small molecules from d). At least two of the compounds seems to increase the myelination of the nanofiber.

To establish our screening assay (Figure S5), PDGFRa-tdTomato+ OPCs from d75-d85 differentiation cultures were purified by magnetic-activated cell sorting (MACS) with thy1.2 microbeads and then plated into 384-well plates. We noticed a small degree of well-to-well variation in the day 0 Nluc activity (relative light unit (RLU) values), which was likely caused by small differences in the number of cells plated per well (Figure S4c).

To control for this variability, instead of using absolute RLU to compare effects of compounds on the cells, we used fold change (FC), which was calculated for each well by dividing a “time-point” RLU of a well by day 0 RLU of the same well (Figures 2a-c). The FC value had better z’ factor values (−0.27 in FC vs −0.37 in raw RLU at d10 time-points) and better relative standard deviation (16% in FC vs 29% in raw RLU at d10 time-point) while retaining the significance of the results (Figure 2a-b). Therefore, RLU fold change values calculated at day 5, 10 and 15 (Figure 2c) were used as readouts for our assay.

The screening assay was validated using a number of previously reported OPC differentiation promoting compounds, and the majority of them showed a dose dependent effect on MBP-secNluc expression (Figure 2d). We have also established an *in vitro* myelination system where purified hOPCs are plated on electrospun nanofibers. The plated OPCs align their processed to the nanofibers as early as two days after plating, and within three weeks they mature into OLs and often myelinate the fibers as observed by PLP1-GFP+ processes (Figure S6) or by immunostaining for MBP protein (*13*). An increased number of PLP1-GFP+ processes aligning and potentially myelinating the nanofibers, within 10 days in culture, was observed only in the presence of a few compounds, such as Tasin-1 and Quetiapine (Figure 2e).

### High Throughput Screen identifies compounds that increase OL maturation

Using the above-described PT-P1-MsNL-based assay system, we screened the Library of Pharmacologically Active Compounds (LOPAC) and TOCRIS small molecule libraries (Tocriscreen Plus) (~2500 compounds total). For an initial screen, two doses (250 nM and 1.25 uM) of the LOPAC compounds and three doses (100 nM, 500 nM, 2.5 μM) of the TOCRIS compounds were tested (Figures 3a-b). A total of 240 compounds (120 from each library) that showed FC increases in secNLuc activity that were greater than two standard deviations above the DMSO vehicle control wells were retested at 7 duplicate doses (15 nM – 11 μM), each at three time-points (5,10 and 15 days) (Figure 3). Since the two screening were run several months apart, the results from each library are reported separately.

**Figure 3.**
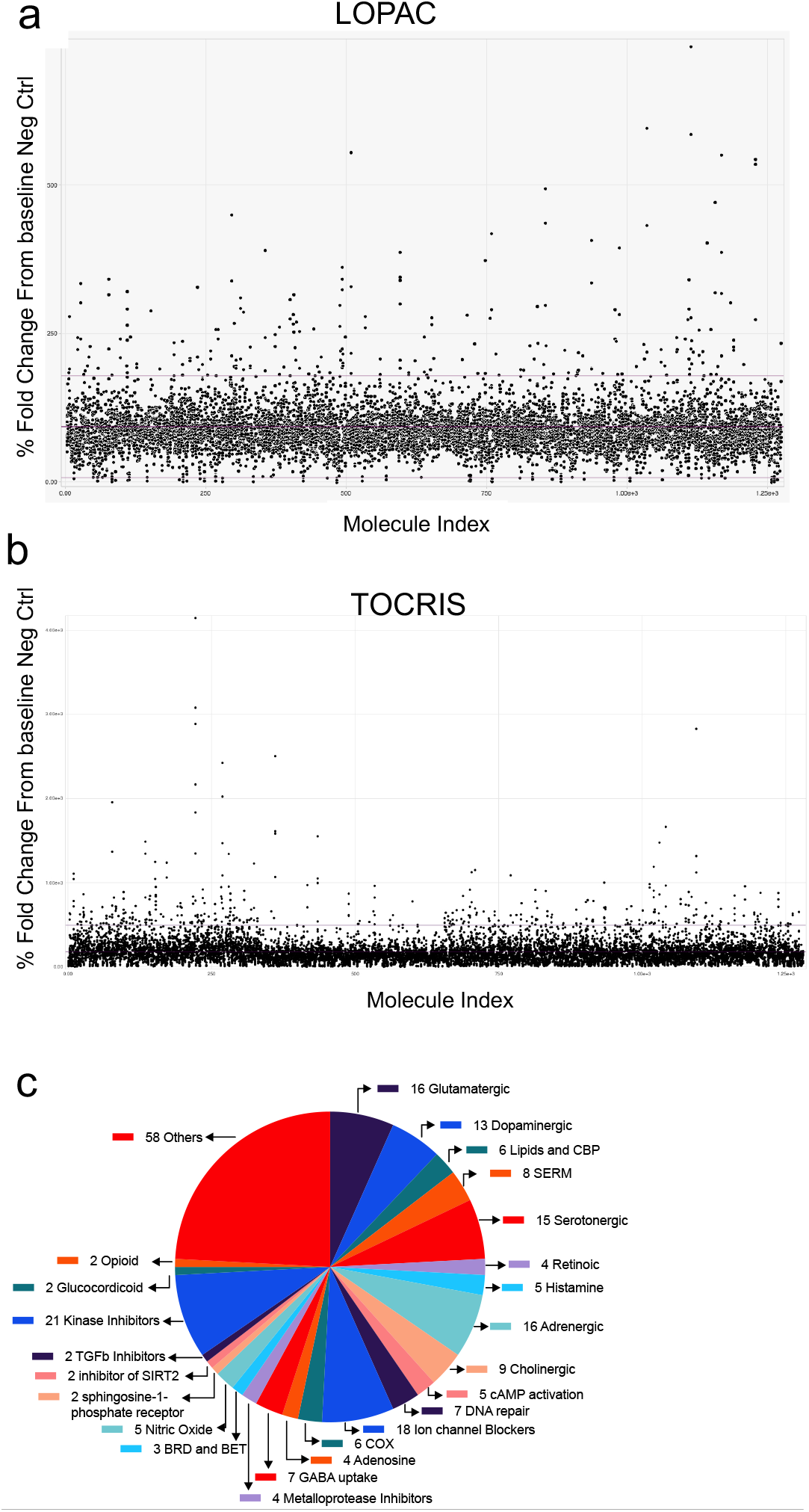
High Throughput Screen for compounds that enhance oligodendrocyte differentiation. **a-b)** Scatterplot of the HTS for compounds that increase activity of MBP-driven secNLuc in the culture media. Plot represents fold change of secNLuc activity (y-axis) versus the compound ID (x-axis) for **a)** LOPAC and **b)** TOCRIS library of small molecules. **c)** A total of 240 compounds with some effect to increase MBP-NanoLuc expression are graphed as a pie-chart according to a class they fall on. The numbers indicate the number of molecules that fall on that class.

The majority of the hit compounds identified in our HTS have known targets such as dopaminergic, adrenergic, cholinergic, cox, DNA repair, GABA uptake, opioid, P450/cholesterol biosynthesis, and SERM (Figure 3c, table 1) that have been identified from previous screens using rodent OPCs. We also identified a number of targets that have been implicated but not well studied in OL biology such as Butyryl cholinesterase(*15*–*17*) (*Tetraisopropyl pyrophosphoramide*), BET bromodomain inhibitor(*18*, *19*) (*I-BET151 and SGC-CBP30*), and NMDA receptor blocker (Ro 25-6981 maleate)(*20*–*22*). We also identified at least two potent compounds that have not been previously reported to promote OL maturation or myelination.

**Table 1.**
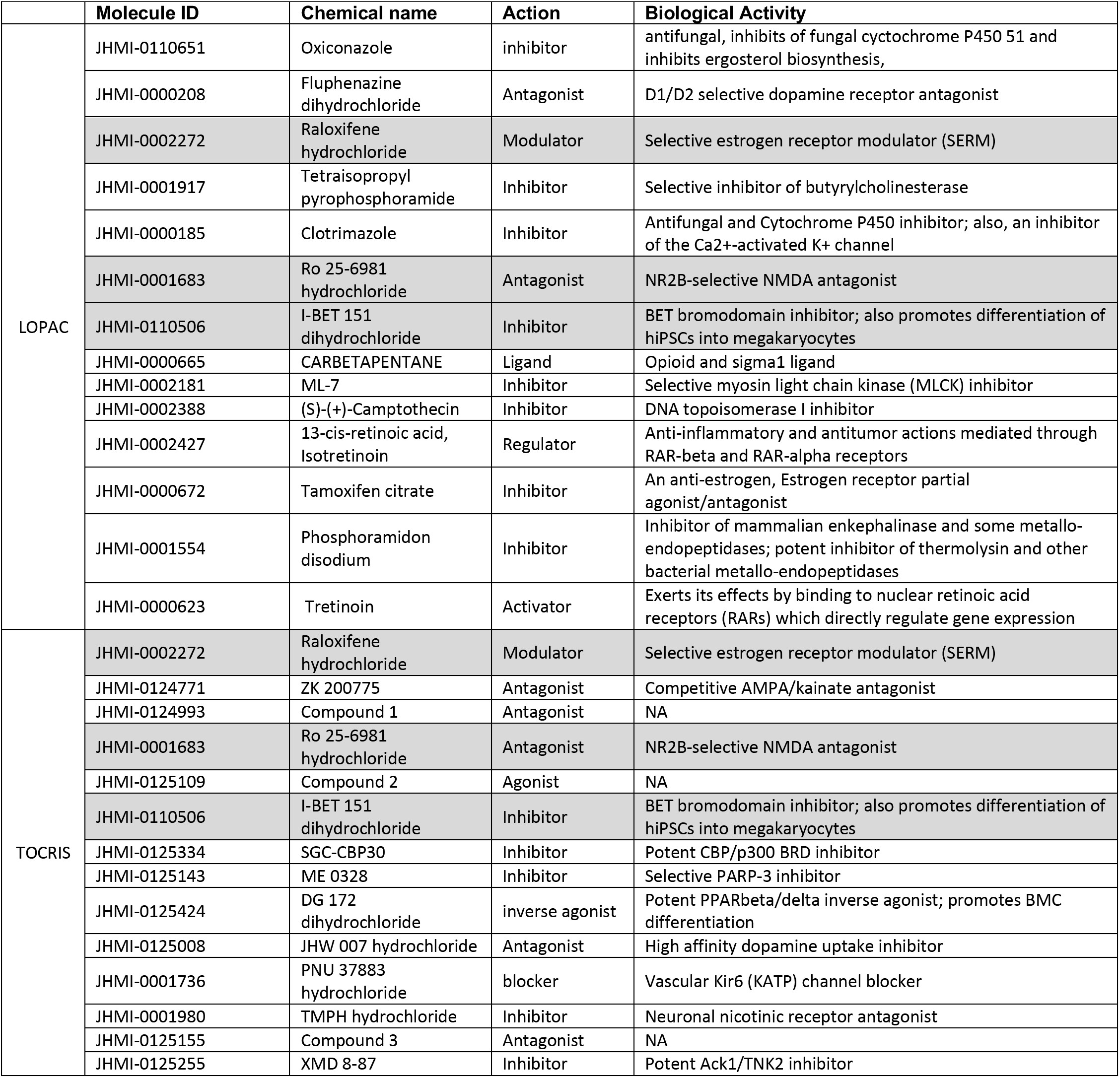
List of top 15 hit compounds from each library. Compounds that are listed in both the libraries are highlighted gray.

### Validation of the promising hit compounds

To validate the Nluc-based screening result that our potential hit compounds indeed enhance OPC maturation to OLs in the human cell system, we performed immunohistochemistry (IHC) analysis of using antibody against MBP and find higher number of MBP+ cells in the hOPCs treated with the two most promising small molecules identified from our screen (Figure 4). Furthermore, in our *in vitro* myelination assay, hOPCs treated with the two potential small molecules or Ro25-6981 for 10 days align their processes to the nanofibers and appear to myelinate them better than DMSO treated cells (Figure 4, lower panel).

**Figure 4.**
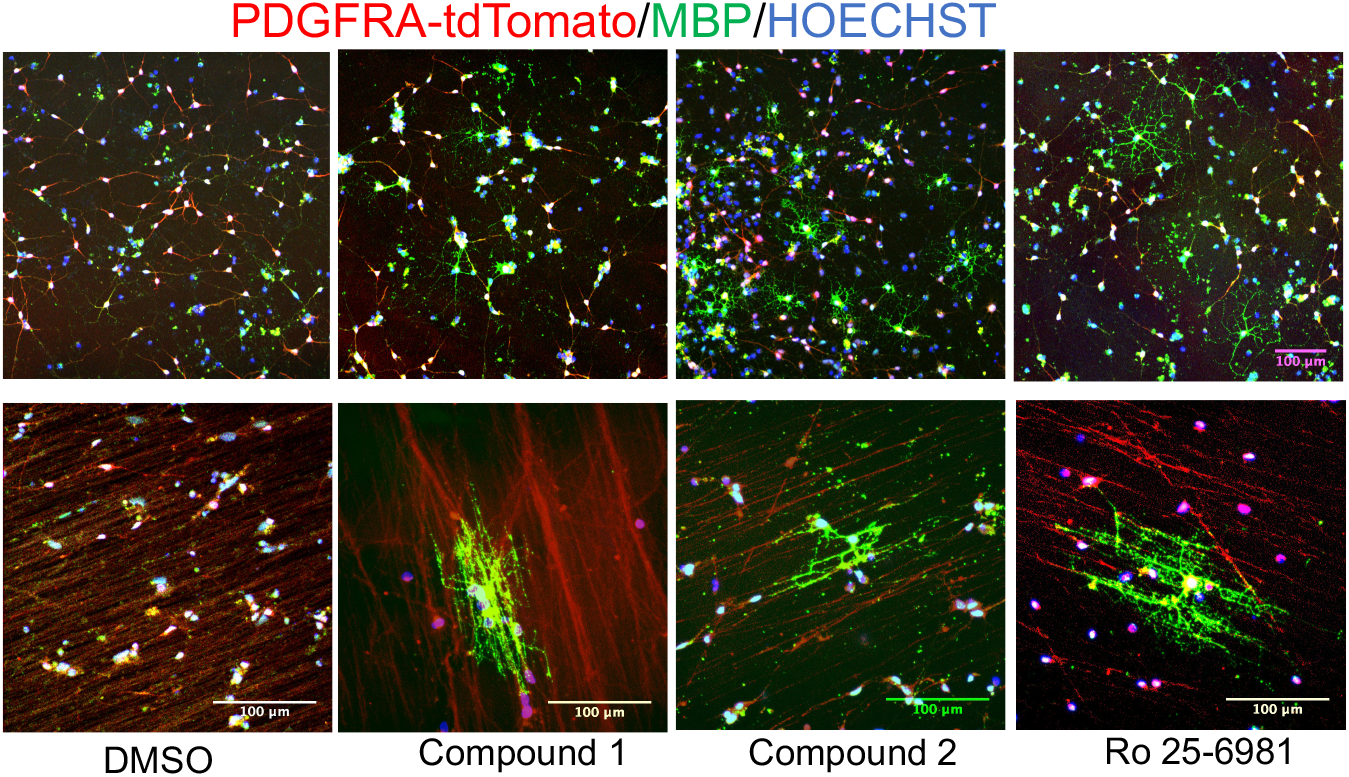
Validation of promising hit compounds. Immunostaining of purified OPCs treated with different compounds for 10 days show stronger MBP staining and more MBP+ cells (top panel) and appearance of improved myelination of electrospun nanofibers ((950 nM diameter nanofibers) lower panel) compared to the DMSO treated cells. Red channel was overexposed to visualize the nanofibers in the lower panel. Immunostaining was independently repeated three times with similar results.

In order to further validate the compounds that we identified using our triple reporter system, we generated another dual reporter system (PDGFRα-P2A-tdTomato-P2A-Thy1.2 and MBP-P2A-secNluc) in an independent, male hESC line (RUES1) and named it RPD-MsNL (Figure 5). MACS-purified PDGFRα-tdTomato expressing OPCs generated from the RPD-MsNL hESC reporter line express other OLC markers and also produce secreted nano-luciferase as expected (Figure 5a-b). Next, we cultured the RPD-MsNL-derived OPCs in the presence of the two most promising hit compounds and Ro25-6981 and validated the efficacy of those compounds in this independent cell line via the Nluc activity, IHC and nanofiber-based myelination assays (Figures 5c-e).

**Figure 5.**
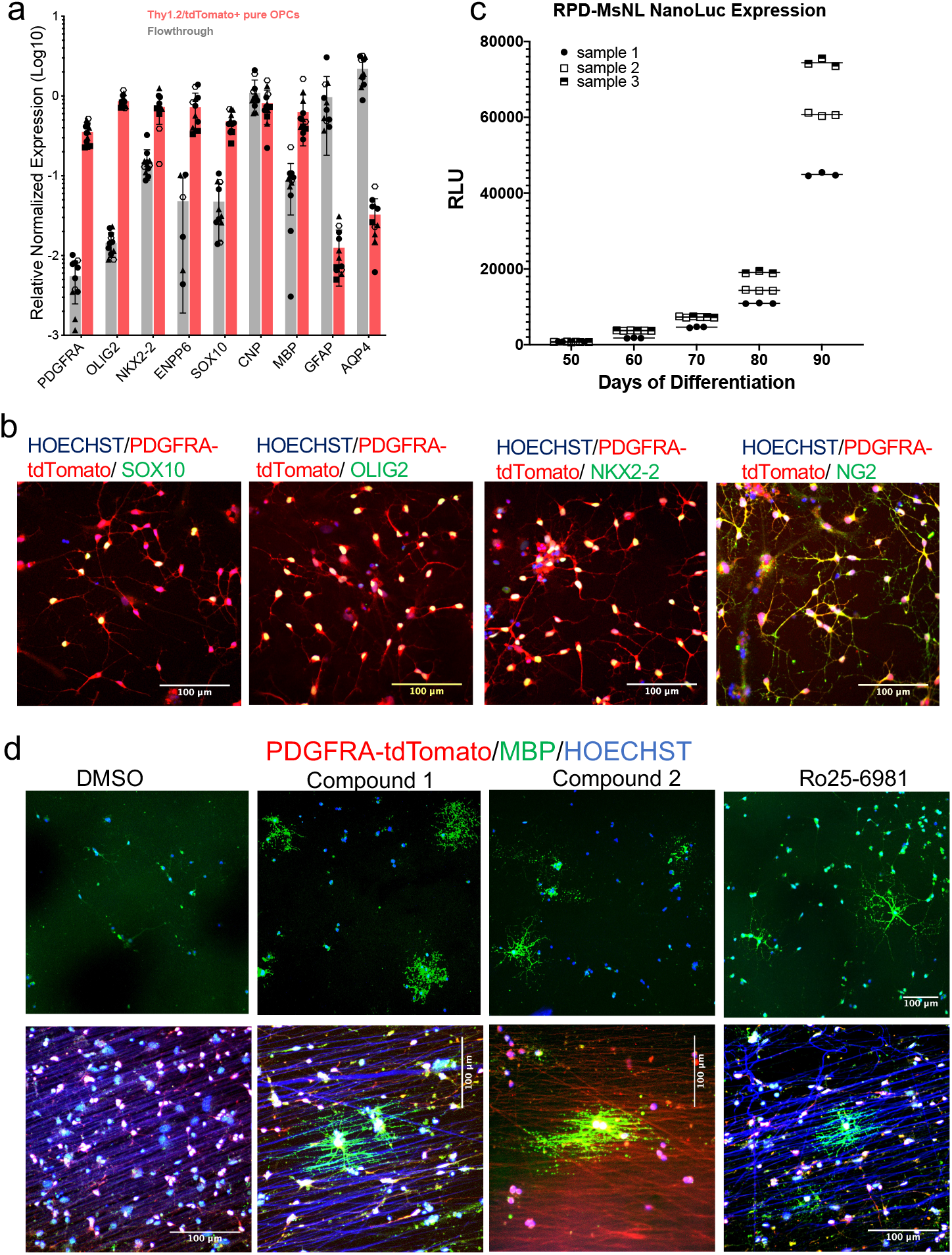
Validation of reporter OPCs generated using an independent, male hESC line (RUES1). **a)** Differentiated OPCs were MACS purified for the expression of PDGFRA-tdTomato-thy1.2, and the expression of different OPC markers between the tdTomato+ and tdTomato-(flow through) population were quantified by qPCR analysis, which shows enrichment of OPC markers in the tdTomato enriched population compared to the flow through. Three biological and three technical replicates each were used for qPCR analysis. Data are presented as mean ± SD. Biological and technical replicates are distinguished by filled vs clear symbols used for each data point. Source data for the qPCR are provided as a Source Data file. **b**) Immunohistochemistry demonstrating that the MACS purified tdTomato+ cells express the OPC markers SOX10, OLIG2, NKX2.2 and NG2. Immunohistochemistry was independently repeated three times with similar results. **c)** secNluc activity in the cell culture medium of differentiating RPD-MsNL reporter line measured with Nano-Glo assay. 20 uL of culture media from different days of differentiation (x-axis) were removed for the assay. Increase in Nano-Luc activity(y-axis), which corresponds to MBP expression increases as cells mature. **d)** Immunostaining of the purified RPD-MsNL OPCs treated with different compounds for 10 days. Stronger MBP staining, more MBP+ cells (top panel) and better myelination of the electrospun nanofibers ((950 nM diameter nanofibers) lower panel) by the compound treated OPCs compared to the DMSO treated cells is noticeable.

## DISCUSSION

Promoting remyelination of neurons in the CNS is a promising approach for treatment of MS and other demyelinating neurodegenerations. If endogenous OPCs could be stimulated to differentiate and remyelinate affected axons, we could reduce and potentially stop disease progression, including the neurodegeneration aspects of the diseases, and might even be able to reverse already established disability in MS patients. Remyelination-based therapy would complement current immune-modulation-based treatments. Therefore, identification of lead small molecules that can promote differentiation of myelinogenic OPCs can have direct translational implications. The systems being used currently to screen for drugs that promote oligodendrocyte maturation and myelin production are predominantly using rodent cells. For developing effective remyelinationbased therapies, validating and expanding the mouse studies to a human system is essential. In addition, drug discovery performed with human cells has a better likelihood to identify leads that will translate into effective treatments. However, a verified human OPCs/OLs-based drug discovery platform has not yet been reported, probably due, in large part, due to the difficulty and challenges in obtaining a large quantity of pure primary glial cells for such work. Since the human stem cell-derived OPCs and OL now provide a powerful and versatile source of human cell culture systems for drug discovery, we set forth to establish a drug screening platform that uses hPSC-derived OPCs and OLs.

We engineered an advanced human embryonic stem cell-based reporter system where florescent proteins and secreted Nano luciferase (secNluc) protein are driven by OPC and OL specific promoters, and used OPCs derived from the reporter hESCs to establish a drug screening platform that allows for high throughput screening of myelination promoting drugs. With the secreted Nluc system, cells don’t need to sacrificed for analsyis, allowing time-course studies and significant cost and research benefits. Using this system, we screened ~2500 bioactive molecules in at least two doses and three time-points. 240 potential hit compounds identified from an initial screen were further tested in duplicates at 7 doses and three-time points. The extensive data on the effect of these compounds on MBP protein expression in human OLs would not have been possible without the reporter system described here. The dose response dataset that we have generated for the ~2500 compounds will be a useful resource for ongoing and future remyelination studies. Furthermore, the reporter cells system that we have established is suitable for a larger scale, human cell-based screen of remyelination compounds.

As a validation of our screening platform, several of the compounds we identified, including muscarinic receptor antagonists (benzotropine, clemastine and carbetapentane) cytochrome P450 inhibitors (clotrimazole, oxiconazole), SERMs (raloxifene, tamoxifen), dopamine receptor antagonists (fluphenazine), and ROCK inhibitors (ML9, ML7) (Table 1) were recently reported as promoting remyelination of rodent OPCs (*9*, *11*, *23*–*26*). In addition, inhibition of bromodomain containing proteins has been reported to help improve myelination in a murine system(*18*, *19*) but its role in human system has not been examined. Tetraisopropyl pyrophosphoramide, an inhibitor of butyrylcholinesterase (BChE), is another compound of interest that was identified in our screen. Additionally, a role of BChE in modulating neuroinflammation, demyelination and neuropathology in MS has been suggested (*27*) (*17*, *28*, *29*), and an inhibitor of BChE, rivastigmine, has been shown to suppress neuroinflammation in EAE models(*29*). Our study provides validation of these targets in a human OPC and OL system.

Possibly due to the use of the highly sensitive Nluc reporter system and also because we used human OPCs for our screens, given that it is well known that human and rodent cells differ in many important ways, we were able to identify a number of targets that have not been previously reported. We show that at least two of these new compounds identified from our screen increase OL maturation in the OPCs derived from two different hESC lines. Using the *in vitro* myelination assay that we have established, we found that these compounds could potentially promote myelination as well. In summary, we have established a robust human cell-based drug screening platform for the discovery of promyelinating compounds, performed a small-scale proof-of-principle drug screen, and identified a number of promising compounds. As a next step, genetic validation of the pathways targeted by these small molecules and *in vivo* testing of the molecules in rodent models of demyelination will be performed to further understand the activity of these molecules, define their target pathways in promoting remyelination, and explore their therapeutic potential.

## ACKNOWLEDGEMENTS

This work was supported by grants from the Gilbert Family Foundation, NIH (P30 EY001765 and K99 EY029011), Sanofi Inc., unrestricted funds from Research to Prevent Blindness, and generous gifts from the Guerrieri Family Foundation.

## AUTHOR CONTRIBUITIONS

W.L. assisted with experimental designs, conducted experiments, conducted bioinformatic analysis, and edited the manuscript. C.B assisted with experimental design and high-throughput screening assay and analysis. Y.H, G.S, and S.M, J.C.D assisted with the screening and RNA work. F.T, X.C, and Y.D assisted in generating the stem cell reporters and differentiation of hEC-derived OPCs. W.F., C.C., H.J., assisted with bioinformatic analysis. D.J.Z designed the study, edited the manuscript, and provided funding support. X.C. designed the study, performed experiments, wrote the manuscript, and provided funding support.

## METHODS

### Human pluripotent stem cells (PSCs) and culture conditions

hESC WA09 and RUES1 (both WiCell), NIH-approved hESC lines (NIH approval number: NIHhESC-10-0062, and 0012), were used for this study. hESCs were maintained in mTesr or mTesr plus (85850 or 100-0276, Stem Cell Technologies) on growth factor-reduced Matrigel (354230, Corning) coated plates at 37C, 10% CO_2_/5% O_2_. hPSC colonies were passaged by dissociating with Accutase (A6964, Sigma-Aldrich). Cells were maintained in stem cell media containing 5 mM blebbistatin (B0560, Sigma-Aldrich) for the first 24 hours after passaging, to improve single cell survival.

Karyotype analysis was performed using a qPCR based hPSC Genetic Analysis Kit (StemCell Technologies, #07550) and KaryoStat Karyotyping (ThermoFisher). Chromosomal Cells were routinely tested for mycoplasma contamination (MycoAlert, Lonza) and only the cells free of contamination were used for OPC differentiation.

### Cloning

A guide sequence targeting the stop codon of the *MBP and PLP1* locus was designed in Deskgen.com. Guide sequence with minimal off target and very high activity score was chosen and cloned into the BbsI restriction site of the Cas9 plasmid (Cas9-P2A-Puro modified from Addgene #62988(*30*)). pSpCas9(BB)-2A-Puro (PX459) V2.0 was a gift from Feng Zhang (Addgene plasmid # 62988; http://n2t.net/addgene:62988). To clone the donor plasmid, a ~2 kb PCR product was amplified from genomic DNA extracted from H9 ES cells and cloned into Zero Blunt TOPO cloning vector (ThermoFisher Scientific). The tdTomato-P2A-Thy1.2 reporter DNA sequence was then introduced into the TOPO-based donor plasmid, precisely upstream of the *MBP* or *PLP1* stop codon using Gibson assembly (New England Biolabs). NanoLuc plasmids was purchased from Promega.

### Generation of Reporter Cell Lines

Method for gene editing and report cell line generation, previously reported by our lab(*13*, *14*, *30*), was followed. Cells were transfected using the Lipofectamine Stem (STEM00001, ThermoFisher Scientific) transfection reagent following the manufacture’s recommended protocol. 0.35 μg Cas9 plasmid containing a gRNA sequence and 0.75 μg of donor plasmid were used for transfection. Two days after the transfection, the cells were selected with 750 ng/mL of puromycin for 24 hours, washed and cultured for 5 more days in the same dish with daily media change. After 5 days, the surviving cells were passaged at 500–1000 single cells per well of a 6 well plate for picking individual colony and PCR-based genotyping (*30*). PCR was performed using the Phusion Flash mastermix (ThermoFisher Scientific) and a 2-step PCR protocol following the manufacturer’s instruction. Clones with clear homozygous knock-In were selected for the MBP-secNluc (Figures S1d, e), and a clone with heterozygous KI was chosen for the For PLP1-sfGFP reporter. For the pLP1-sfG KI clone, another allele of the PLP1 locus was sequenced to make sure that it had no mutation.

### Oligodendrocyte Differentiation Protocol

Previously published and well-established oligodendrocyte differentiation protocol (*31*) was followed. Briefly, hESCs were dissociated to single cells and plated on Matrigel coated plate at 100,000 cells/well of a 6-well plate and maintained in mTesr plus at 37C, 10% CO2/ 5% O2. For RUES1 cells 200K cells were used to start differentiation. Two days after passaging, neural differentiation and spinal cord patterning was induced through dual SMAD inhibition (SB431542, 10uM and LDN193189, 250 nM) and 100 nM all-trans RA (*32*). From day 8 to day 12, differentiating cells were maintained in neural induction media supplemented with RA (100nM) and SAG (1 mM). At day 12, adherent cells were lifted and cultured in low-attachment plates to favor sphere aggregation. At day 30, spheres were plated into poly-L-ornithine/laminin-coated dishes in a media supplemented with, B27 (Thermo Fisher, 12587010), N2 supplement (Thermo Fisher, 17502048), PDGF-AA (221-AA-10, R&D systems), neurotrophin-3, HGF (294-HG-025 R&D systems), and T3.

### Flow Cytometry and MACS Purification of the Reporter hOPCs

For flow cytometry analysis and MACS purification, cells were dissociated into single cell suspension by incubating in accutase for ~45 minutes. The single cell suspension was then passed through a ~70 uM cell strainer (BD Biosciences), washed, and resuspended in Live Cell Imaging Solution (ThermoFisher Scientific) for flow analysis or the MACS buffer (Miltenyi Biotec (Auburn, CA)) for MACS cell sorting. Flow analysis was performed with an SH800S Cell Sorter (Sony Biotechnology, San Jose, CA). BSC and FSC was used to select and subset live cells, and only live cells were used to quantify number of tdTomato+ or GFP+ cells. A gate was setup using WT hES cells differentiated to day 95. MACS purification was performed by following manufacturer’s instructions with minor modifications. Cells were resuspended in MACS buffer after passing through cell strainer. A CD90.2 (THY1.2) or O4 MicroBeads were added to the single cell suspension that is resuspended in MACS buffer and incubated at room temperature for 15 minutes for cell binding. Cells were run through the LS or MS magnetic column, columns were washed 3 times with the MACS buffer, and the cells bound to the column were recovered by pushing the cells through using a syringe provided. To increase the purity of tdTomato+ population, the cell collected from the column were run through a new column without additional supplementation of MicroBeads.

### Nanofiber-based myelination assay

Aligned polycaprolactone (PCL) nanofibers was electrospun on a glass coverslip according to a previously reported method (*33*, *34*). The average diameter of the fibers collected was 950 nm ± 200 nm. The nanofibers on coverslips were sterilized by soaking in 95% ethanol, and then washed three times with PBS. The nanofiber coverslips were placed a 24 well plate and coated with PLO (50μg/mL) for 4 hours followed by laminin (10μg/mL) at 37°C overnight. Before cell seeding, laminin was aspirated and the coverslip was allowed to dry completely. For each well of a 24 well plate containing a nanofiber coverslip, 5000 purified OPCs were resuspended in 50 ul of PDGF media and the cells were added to the center of the coverslip. Cells were allowed to attach for 30 mins before gently adding 500 uL of the PDGF media/well to culture the cells. Media with small molecules or DMSO was replenished every 4-5 days. At the end of the experiment. media was removed, cells were fixed with 4% PFA, and used for immunofluoresce staining and imaging.

### Immunofluorescence Staining, Microscopy, and qRT-PCR

Total RNA was isolated using the RNeasy Mini Kit (QIAGEN) and reverse transcribed using the High-Capacity cDNA Reverse Transcription Kit (Applied Biosystems). A 2uL PCR reaction was setup using acoustic liquid handler (ECHO 550, Labcyte) and performed with the CFX384 real-time PCR instrument (Bio-Rad). Assays included at least two technical and two biologicals replicates and were run using the Sso Advanced Universal SYBR Green Supermix (Bio-Rad).

For immunofluorescence staining, cells were fixed with 4% paraformaldehyde, and simultaneously permeabilized and blocked with 0.2% Triton X-100 + 5% BSA + 5% normal goat serum (or a serum specific to the host of secondary antibody) for an hour. Cells were then incubated with appropriate dilution of primary antibodies over-night followed by secondary antibodies for 2 hours. Fluorescence images were taken using either the EVOS FL Auto 2 (ThermoFisher Scientific) or Zeiss 510 confocal microscope.

### Statistical Analysis

All qRT-PCR data are presented as fold change in RNA normalized to the expression of two housekeeping genes: either *GAPDH* and *SRT72* or *GAPDH* and *CREBBP*. qRT-PCR data was analyzed using CFX Maestro (Biorad) qPCR analysis software and graphed using Prism (GraphPad, V9).

### High-Throughput screening

MACS purified reporter OPCs from day 75-85 differentiating culture was used for screening. 1.5K cells/well in of purified OPCs were plated in PLO-laminin coated 384 well plates using the EL406 (BioTeK) peristaltic liquid dispenser. 50uL of PDGF media was used per well to plate the cells. On the day of drug treatment (on day 2 after plating the cells) 20 uL of culture media was removed from each well using an integra ViaFlow 384, and Nluc activity in the media was measured (day 0 reading).

Nluc activity was measured using NanoGlo luciferase assay reagents (Promega N1150) following manufacture’s instruction. Of note, 2.5 ul NanoGlo reaction mix diluted with 2.5 uL of water (1:1 dilution) was used for the assay. i.e 5 uL of diluted NanoGlo reaction mix was added per well containing 20uL of media. RLU reading was performed with a microplate reader (ClariOstar, BMG Labtech) using the preloaded settings for Nano luciferase and 0.1s exposure.

Drug dispensing was performed using ECHO 555, acoustic liquid handler. First, an intermediate plate with 10 uL of glial media in each well of a new sterile 384 well plate was prepared. Appropriate volume of small molecules was dispensed into each well using the ECHO. Additional 10 uL media was added to each wells containing the small molecules. Using integra ViaFlow 384, 20 uL of the media containing small molecules was transferred from the intermediate plate to the plate containing cells. After adding the small molecules cells were cultured in 37 C, 5% Co2 incubator. Every 5 days, 20 uL of media was removed, Nluc activity measured and fresh media with small molecules was replenished as described above. On day 15, cells were fixed with 4% PFA, washed with PBS and either stored in 4C or immediately imaged using a High-Content Imager (Cellomics CX7).

