## supplemental figures for "High-Throughput Screening for Myelination Promoting Compounds Using Human Stem Cell-derived Oligodendrocyte Progenitor Cells Identifies Novel Targets"

**Supplemental Figure 1. Generation of the PT-P1-MsNL reporter cell line. a)** schematics of the plasmids used to confirm that the secNLuc protein product separated by P2A is functional. A tdTomato-P2A sequence was cloned into the original CMV-secNluc plasmid to generate a CMV-tdTomato-P2A-secNluc plasmid. **b)** Different amount of the commercially available CMV-secNluc plasmid and the CMV-tdTomato-P2A-secNluc plasmid were transfected into the HEK293 cells. **c)** Successful detection of Nluc activity in the culture media of the cells transfected with both plasmids. PCR genotyping **d)** followed by sanger sequencing of both bands **e)** was used to confirm CRISPR-Cas9 mediated successful knock-in of the sfGFP reporter sequence into *PLP1* locus of PD-TT reporter line. **f)** PCR genotyping after P2A-secNluc sequence was knocked-in into the MBP locus of the PD-TT and PLPsfGFP dual reporter to make the final triple reporter PT-P1-MsNL cell line.

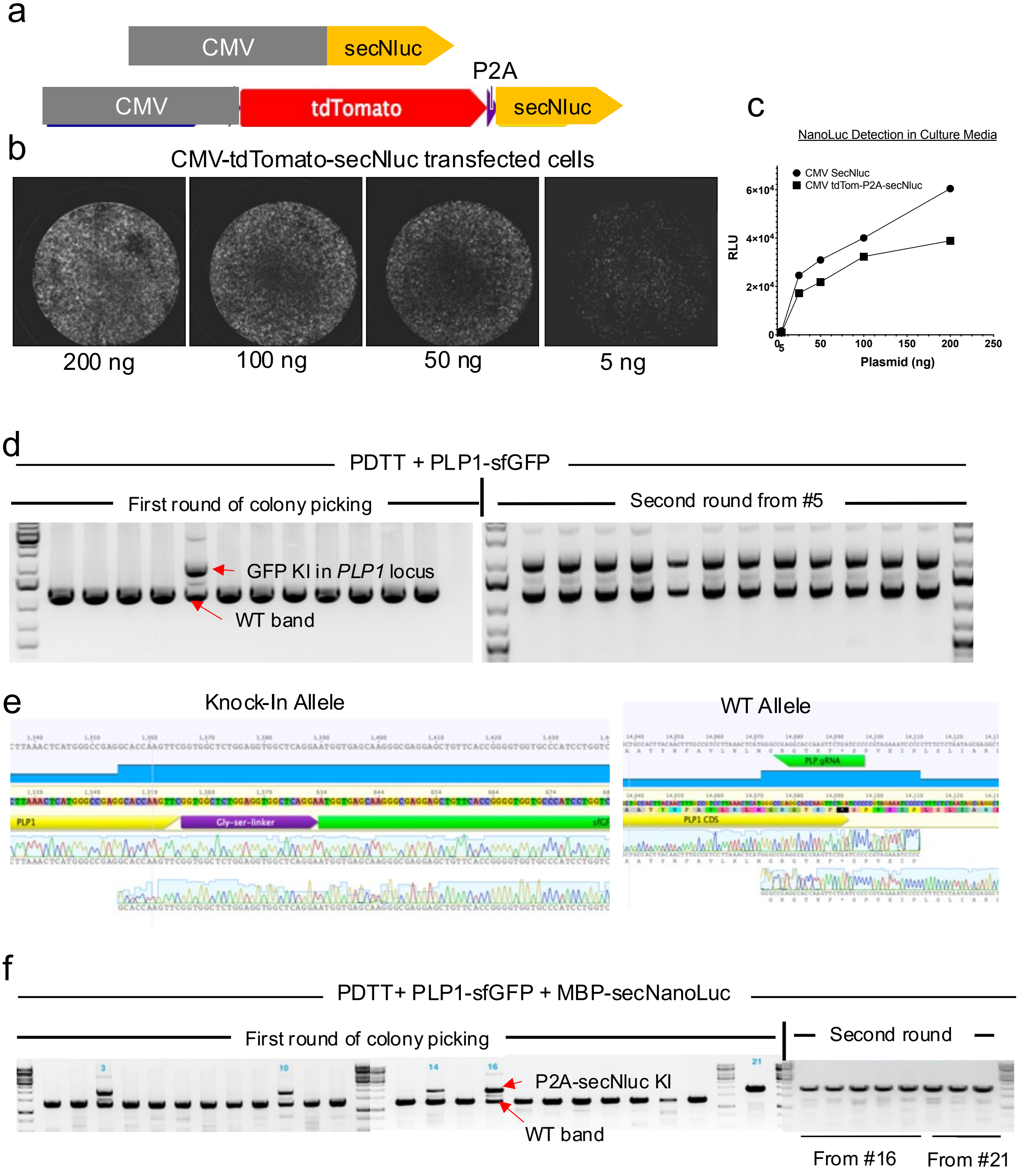

**Supplemental Figure 2**. **Differentiation of PtT-P1-MsNL hESC reporter line to oligodendrocyte cells**. **a)** Expression of the tdTomato and sfGFP fluorescent reporters in the differentiating oligodendrocyte cultures. First PLP1-sfGFP expressing cells appear around 60 and increase as the cells mature. tdTomato expressing cells, on the other hand, is reduced as the cells mature.

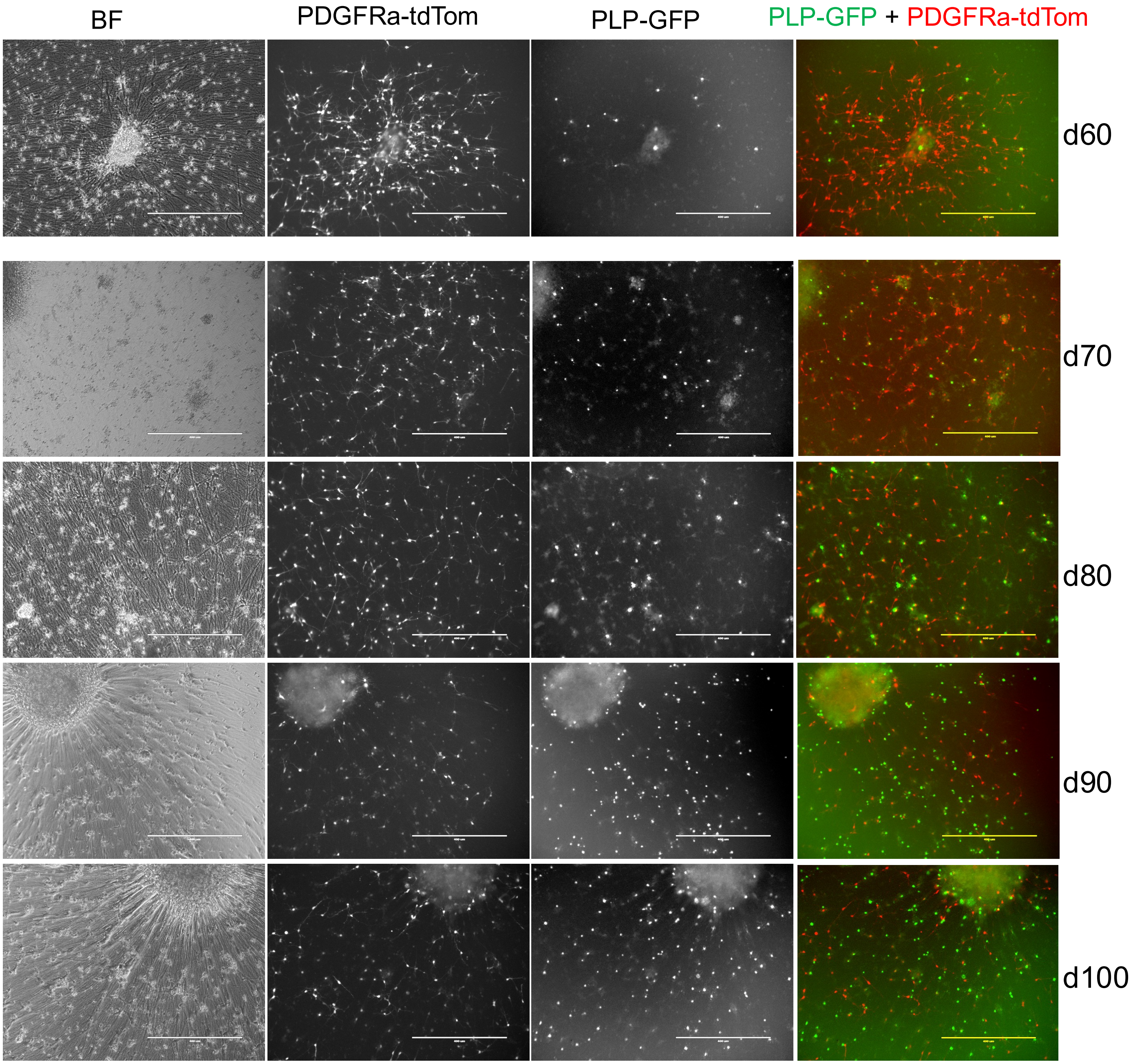

**Supplemental Figure 3. Flow analysis of the reporter OPCs. a)** Gating strategy used to select live cells from the differentiating hOPC population. This BSC and FSC setup was used for all the flow analysis performed. **b)** Flow analysis showing PDGFRa-tdTomato+ and PLP1-sfGFP+ OPCs in the culture differentiated for 90 days. **c)** Significant enrichment of the tdTomao+ as well as GFP+ cells can be achieved by MACS purification using O4 antibody conjugated microbeads.

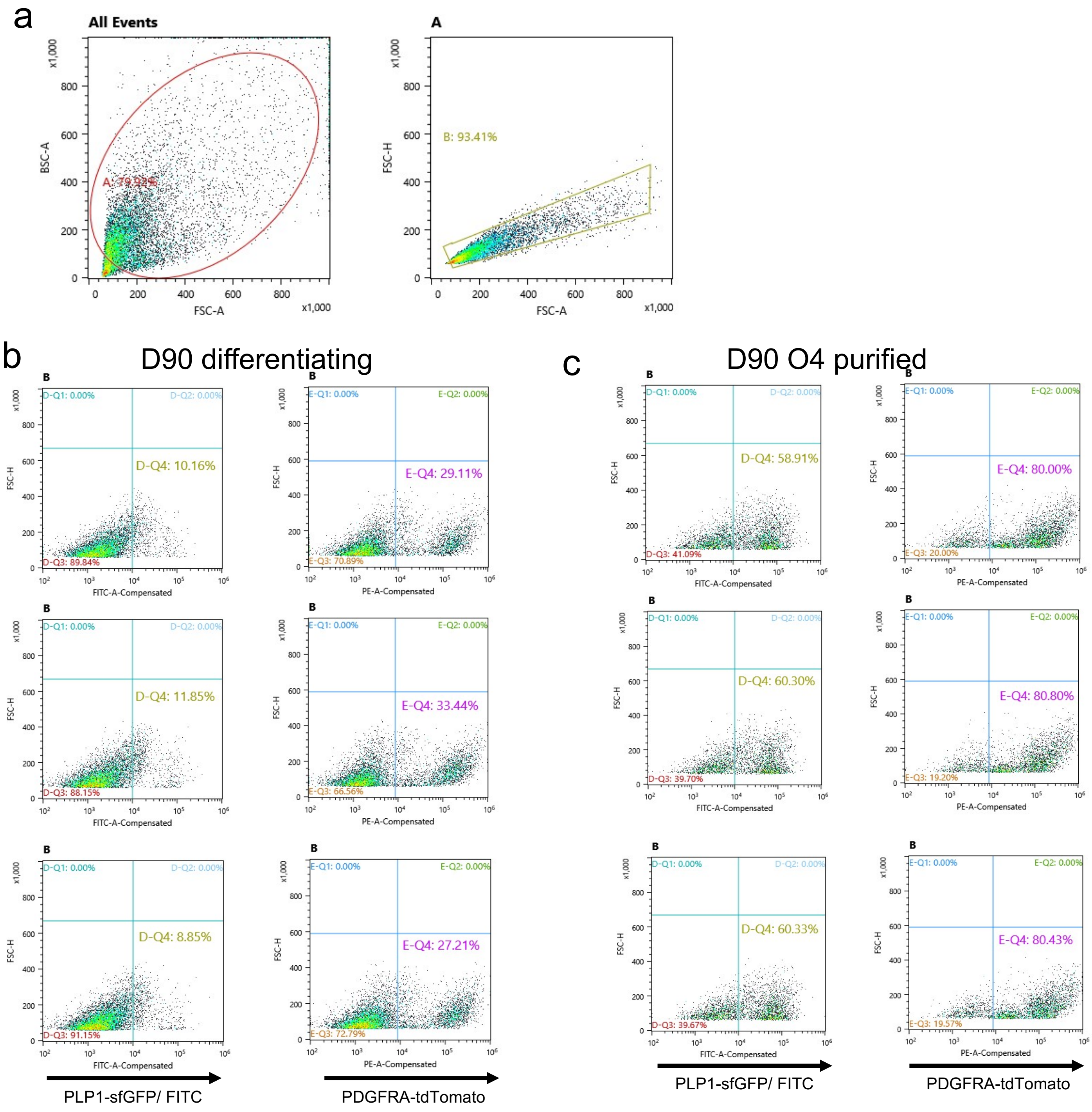

**Supplemental Figure 4. Optimization of the drug screening assay using PT-PG-MsNL hESC-dericed hOCPs. a)** Nluc activity was measured in different volumes cell culture media containing Nluc. **b)** Different volume of NanoGlo reagent was also tested. 20 uL of media and 2.5 uL of Nluc reagent was then used for further optimization. **c)** 1500 cells/well of a 384 well plate (7 plates) was plated to calculate variability caused by initial cell plating. **d)** The OPCs mature and produce more Nluc as a natural course of differentiation, even in control condition (i.e DMSO control). The well-to-well variability caused due to initial cell plating is negligible when compared to the RLU reading of later time-point. **e)** Dose depended effect of of Tasin-1 in MBP-Nluc expression at different time-points. **f**) D10 value for 250 nM Tasin-1 presented as scatterplot to show well to well variability.

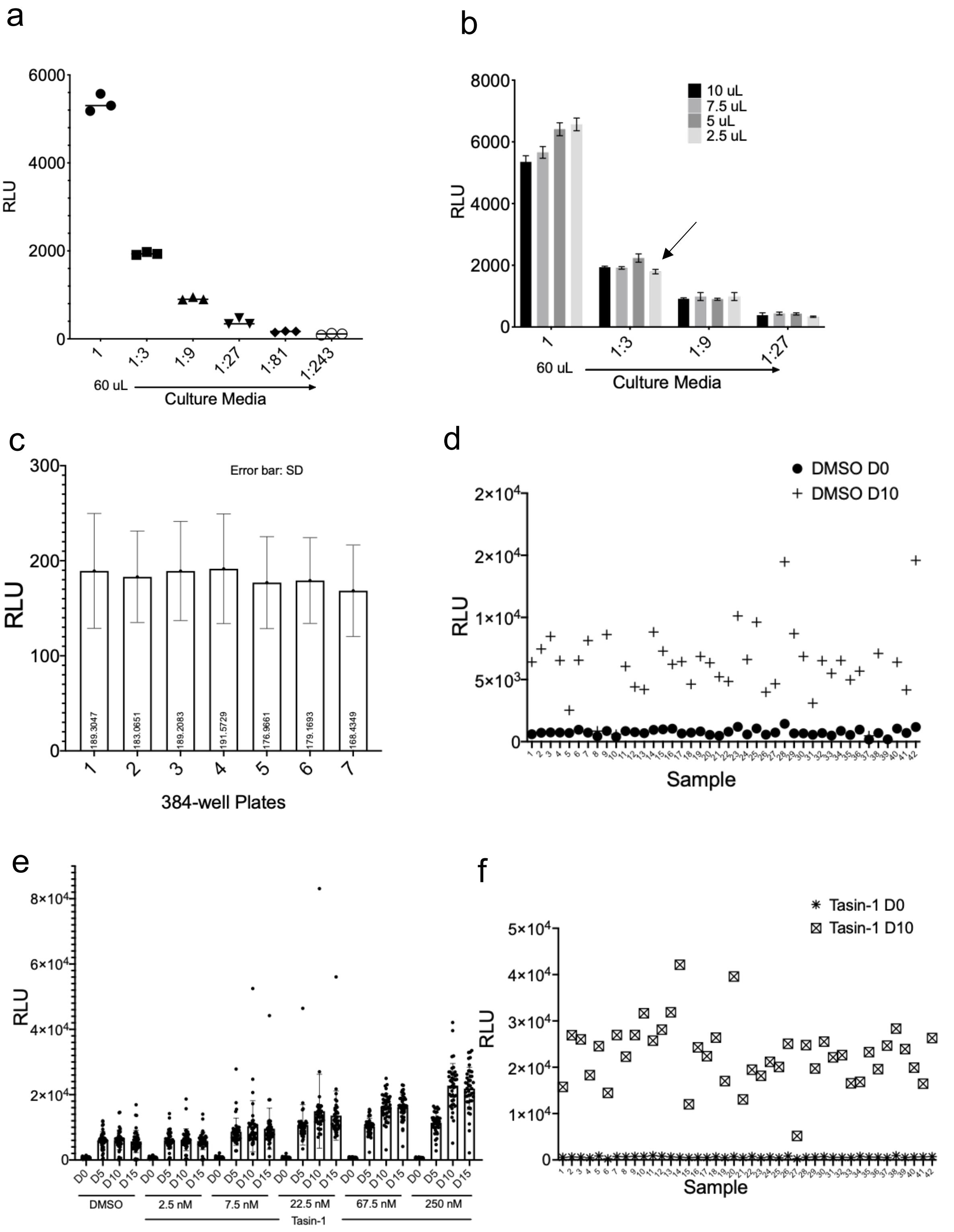

**Supplemental Figure 5. Workflow of the drug screening setup.** A schematic presenting simplified version of drug screening workflow established in our laboratory to identify pro-myelinating compounds using human hPSC-derived OPCs.

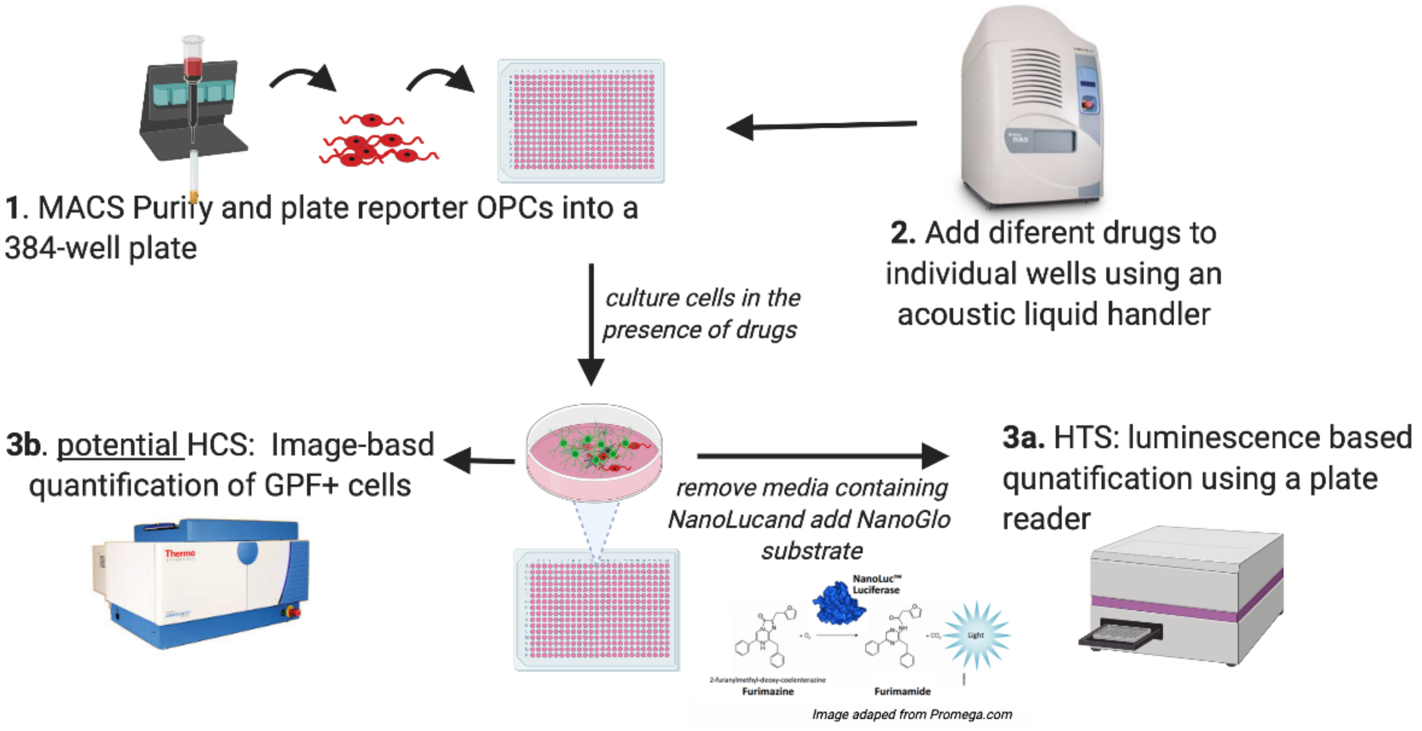

**Supplemental Figure 6**. ***In vitro* myelination culture**. **a)** purified OPCs when plated on a plate containing 950 nM of electrospun nanofibers align their processes within 2 days of plating the cells. **b)** In 3 weeks they mature into OLs and appear to myelinate the fibers.

a PDGFRa-tdTom Nanofibers Merge

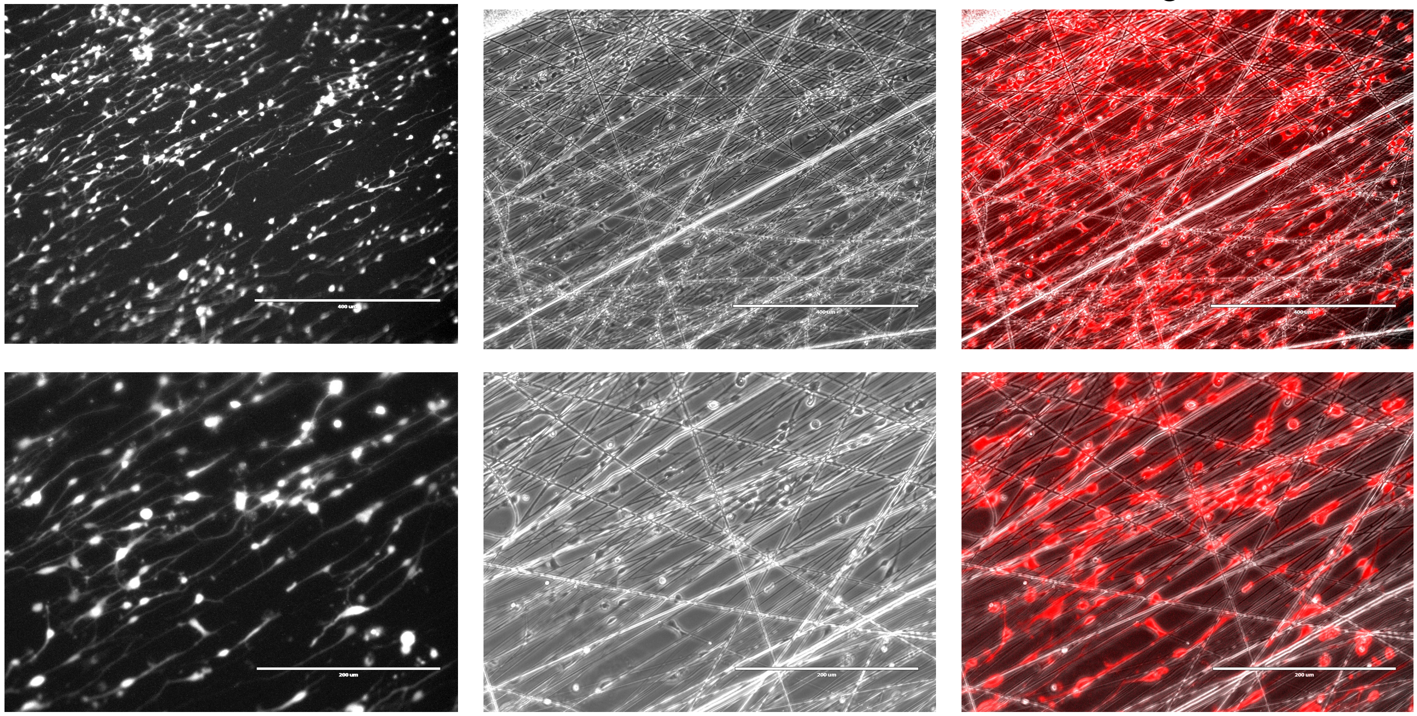

b PLP1-sfG/ HOECHST

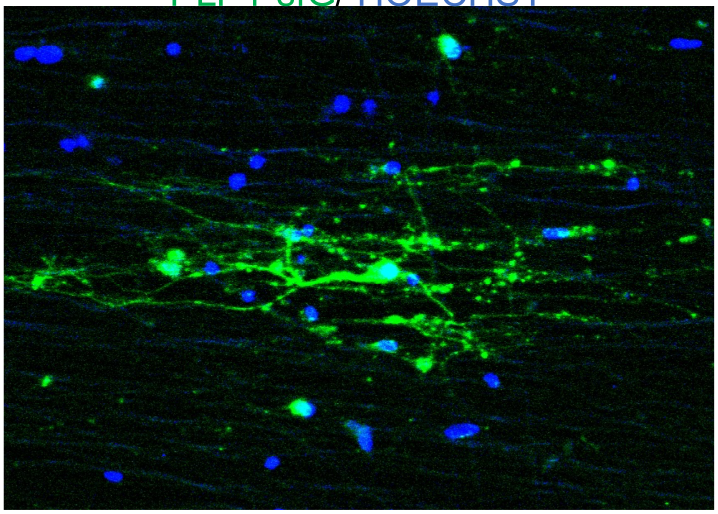

**Supplemental Figure 7. RPD-MsNL reporter cells express OPC markers. a**) Immunohistochemistry demonstrating that the MACS purified tdTomato+ cells express the OPC markers SOX10, OLIG2, NKX2.2 and NG2.

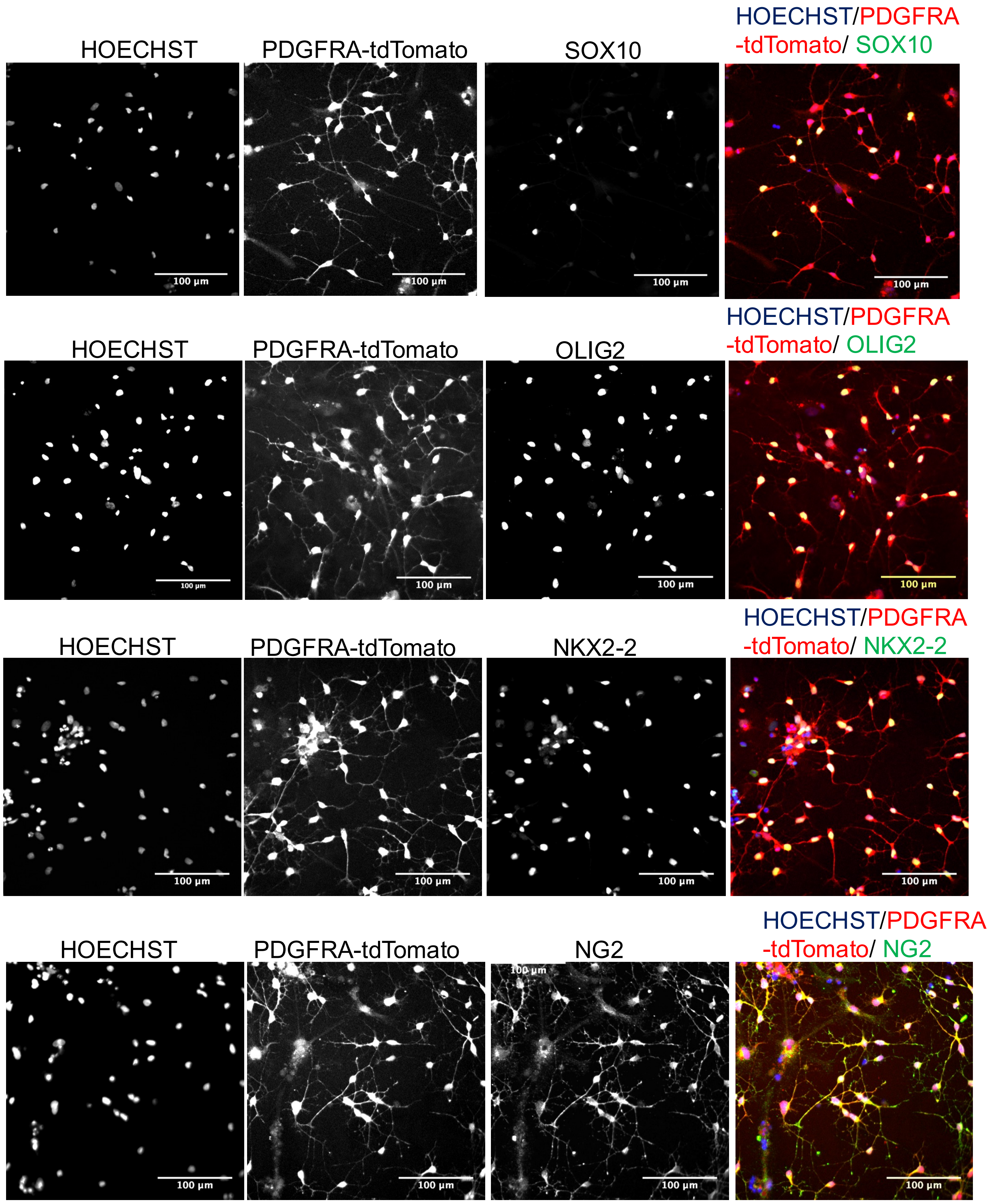
